# Distinct antibody responses to endemic coronaviruses pre- and post-SARS-CoV-2 infection in Kenyan infants and mothers

**DOI:** 10.1101/2022.06.02.493651

**Authors:** Caitlin I. Stoddard, Kevin Sung, Ednah Ojee, Judith Adhiambo, Emily R. Begnel, Jennifer Slyker, Soren Gantt, Frederick A. Matsen, John Kinuthia, Dalton Wamalwa, Julie Overbaugh, Dara A. Lehman

## Abstract

Pre-existing antibodies that bind endemic human coronaviruses (eHCoVs) can cross-react with SARS-CoV-2, the betacoronavirus that causes COVID-19, but whether these responses influence SARS-CoV-2 infection is still under investigation and is particularly understudied in infants. In this study, we measured eHCoV and SARS-CoV-1 IgG antibody titers before and after SARS-CoV-2 seroconversion in a cohort of Kenyan women and their infants. Pre-existing eHCoV antibody binding titers were not consistently associated with SARS-CoV-2 seroconversion in infants or mothers, though we observed a very modest association between pre-existing HCoV-229E antibody levels and lack of SARS-CoV-2 seroconversion in infants. After seroconversion to SARS-CoV-2, antibody binding titers to endemic betacoronaviruses HCoV-OC43 and HCoV-HKU1, and the highly pathogenic betacoronavirus SARS-CoV-1, but not endemic alphacoronaviruses HCoV-229E and HCoV-NL63, increased in mothers. However, eHCoV antibody levels did not increase following SARS-CoV-2 seroconversion in infants, suggesting the increase seen in mothers was not simply due to cross-reactivity to naively generated SARS-CoV-2 antibodies. In contrast, the levels of antibodies that could bind SARS-CoV-1 increased after SARS-CoV-2 seroconversion in both mothers and infants, both of whom are unlikely to have had a prior SARS-CoV-1 infection, supporting prior findings that SARS-CoV-2 responses cross-react with SARS-CoV-1. In summary, we find evidence for increased eHCoV antibody levels following SARS-CoV-2 seroconversion in mothers but not infants, suggesting eHCoV responses can be boosted by SARS-CoV-2 infection when a prior memory response has been established, and that pre-existing cross-reactive antibodies are not strongly associated with SARS-CoV-2 infection risk in mothers or infants.

## Introduction

The SARS-CoV-2 pandemic has caused global catastrophe and is characterized by varying infection risk and clinical outcomes in those that become infected. Younger age has been associated with lower likelihood of infection in numerous studies [1, 2]. Several explanations for this phenomenon have been hypothesized, including the influence of cross-reactive immune responses to endemic human coronaviruses (eHCoVs), also known as seasonal or common-cold causing human coronaviruses. Many studies have shown that eHCoV antibody levels are increased upon SARS-CoV-2 infection [3–11], which may indicate “boosted” pre-existing memory responses that are cross-reactive. It remains unclear whether such cross-reactive antibody responses could modulate SARS-CoV-2 infection risk.

Additionally, while several studies have examined eHCoV antibody responses in children and adults [12], studies testing for eHCoV antibody responses in newborns or infants, and studies that directly compare infants and adults are lacking. Infants are born with passively transferred eHCoV antibodies from their mothers that wane during the early months of life. Those < 6 months of age are less likely to experience eHCoV infection compared to older children [13, 14] and thus will not have memory responses that can be further stimulated by another HCoV infection. In addition, when infants are infected, their antibody responses may differ from those of adults [15, 16], further underscoring the importance of studying eHCoV and SARS-CoV-2 antibody dynamics in infant populations.

Here, we profiled eHCoV antibodies in infants and mothers by measuring IgG titers to the spike protein of four eHCoVs, including two from the same genus as SARS-CoV-2 (betacoronaviruses HCoV-OC43 and HCoV-HKU1), and two alphacoronaviruses (HCoV-229E and HCoV-NL63). We also measured antibodies to the SARS-CoV-1 spike protein, which shares the most sequence homology with SARS-CoV-2 among the coronaviruses we included (76% identity, [17]). We leveraged a longitudinal cohort with mothers and infants that did or did not seroconvert to SARS-CoV-2 to 1) test for differences in eHCoV antibody titers between infants and mothers in naïve and SARS-CoV-2-seroconverted samples, and 2) evaluate associations between pre-existing eHCoV titer and SARS-CoV-2 seroconversion during the study period.

## Results

### Participant groups and longitudinal sample timing

Longitudinal plasma samples collected from an ongoing study of mother-to-child virome transmission in Nairobi, Kenya (the Linda Kizazi cohort) were previously tested for SARS-CoV-2 nucleocapsid seroconversion by enzyme-linked immunosorbent assay (ELISA) (Begnel et al., in revision). Mothers and infants were grouped as either seroconverters or never-seropositive for SARS-CoV-2 during the follow-up included in this sub-study (from April 2019-December 2020; **Figs. 1A and 1B**). Plasma samples from seroconverters (N = 50) included pre-pandemic (as available prior to October 2019; mothers, N = 14; infants, N = 5), last seronegative (mothers, N = 35; infants, N = 11), and first seropositive samples (mothers, N = 36; infants, N = 14). For individuals that never seroconverted in the study period (N = 121), we selected a pre-pandemic sample (when available; mothers N = 21; infants, N = 10), as well as a pandemic-era sample, termed “time matched seronegative”, that overlapped the time period (December 2019-April 2020) of the last negative samples from seroconverters (mothers, N = 62; infants, N = 56; **Figs. 1A and 1B**). None of the mothers seroconverted during pregnancy, so any detectable SARS-CoV-2 antibodies in infant plasma used to determine serostatus (Begnel et al., in revision) were a result of postnatal infection and not due to passive transfer of SARS-CoV-2 antibodies in utero. Median (IQR) infant age in sample groups were as follows: pre-pandemic, 9.7 (6.7-10.6) weeks; last negative or time-matched negative, 25.1 (10.0-38.8) weeks; first seropositive, 47.4 (33.0-65.7) weeks. In the Linda Kizazi cohort, approximately 20% of infants and mothers reported one or more mild to moderate symptoms of COVID-19 at their first seropositive visit or since their last seronegative visit and there were no reported hospitalizations or deaths due to COVID-19 in the cohort (Begnel et al., in revision).

**Figure 1.**
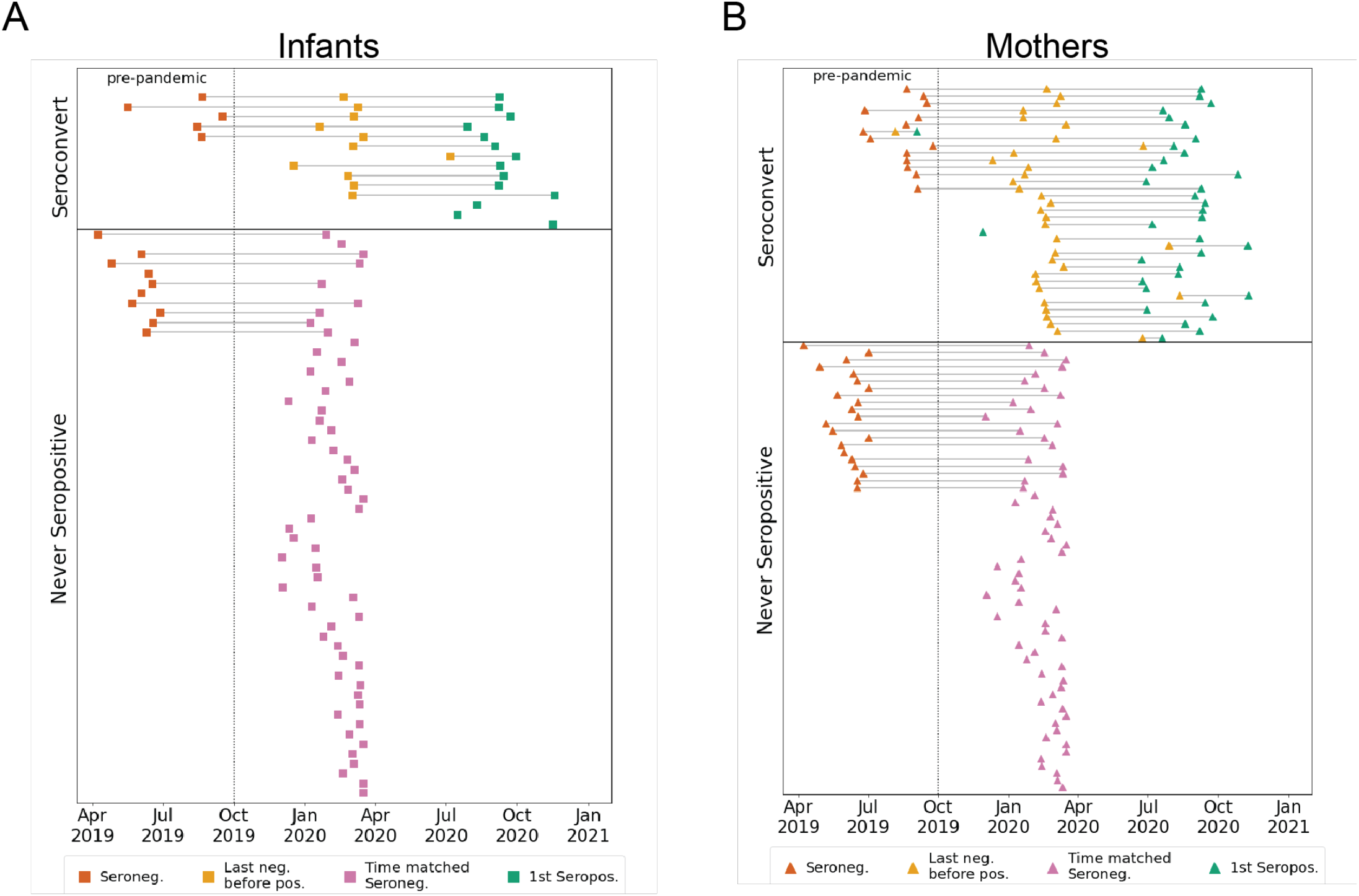
Participant groups and plasma sample timing. (A) Infants or (B) mothers were grouped based on nucleocapsid ELISA results (Begnel et al., in revision) as either seroconverting or never seropositive in the sampling window from April 2019 to December 2020. Samples from seroconverters were selected as pre-pandemic (red), last seronegative (yellow), and first seropositive (green). For never seropositive individuals, pre-pandemic (red), and pandemic-era samples (pink) that overlap the calendar time window of the last seronegative samples in the seroconverting group, were selected.

### Longitudinal eHCoV antibody responses in SARS-CoV-2 seroconverting and non-seroconverting mothers and infants

To test for the presence of IgG antibodies targeting eHCoVs, we compared antibody binding titers to spike from the four commonly circulating eHCoVs in longitudinal plasma samples from infants and mothers using a multiplexed chemiluminescent antibody binding immunoassay. In pre-pandemic samples collected prior to the emergence of SARS-CoV-2, mothers and infants displayed similar levels of antibodies against all four eHCoVs, as would be expected for maternally transferred antibodies present in infants at the age sampled, which was a median of 9.7 weeks of age (**Fig. 2**, left column). In pandemic-era samples collected most recently before SARS-CoV-2 seroconversion or in the time-matched window for non-seroconverting individuals, mothers had significantly higher levels of antibodies targeting the four eHCoVs compared to infants, which were a median of 25.1 weeks of age, with the most pronounced difference for HCoV-NL63 (**Fig. 2**, middle column). This difference was largely driven by lower median levels of infant antibodies, rather than an increase in maternal antibodies, suggesting waning of the passively transferred response in infants over time. Similarly, upon seroconversion to SARS-CoV-2, mothers exhibited significantly higher levels of eHCoV antibodies than infants (**Fig. 2**, right column). These results demonstrate differences in infant and maternal antibody responses to eHCoVs in this cohort likely reflecting the more limited opportunity for eHCoV exposure in the early months of infant life in part due to passive antibody protection.

**Figure 2.**
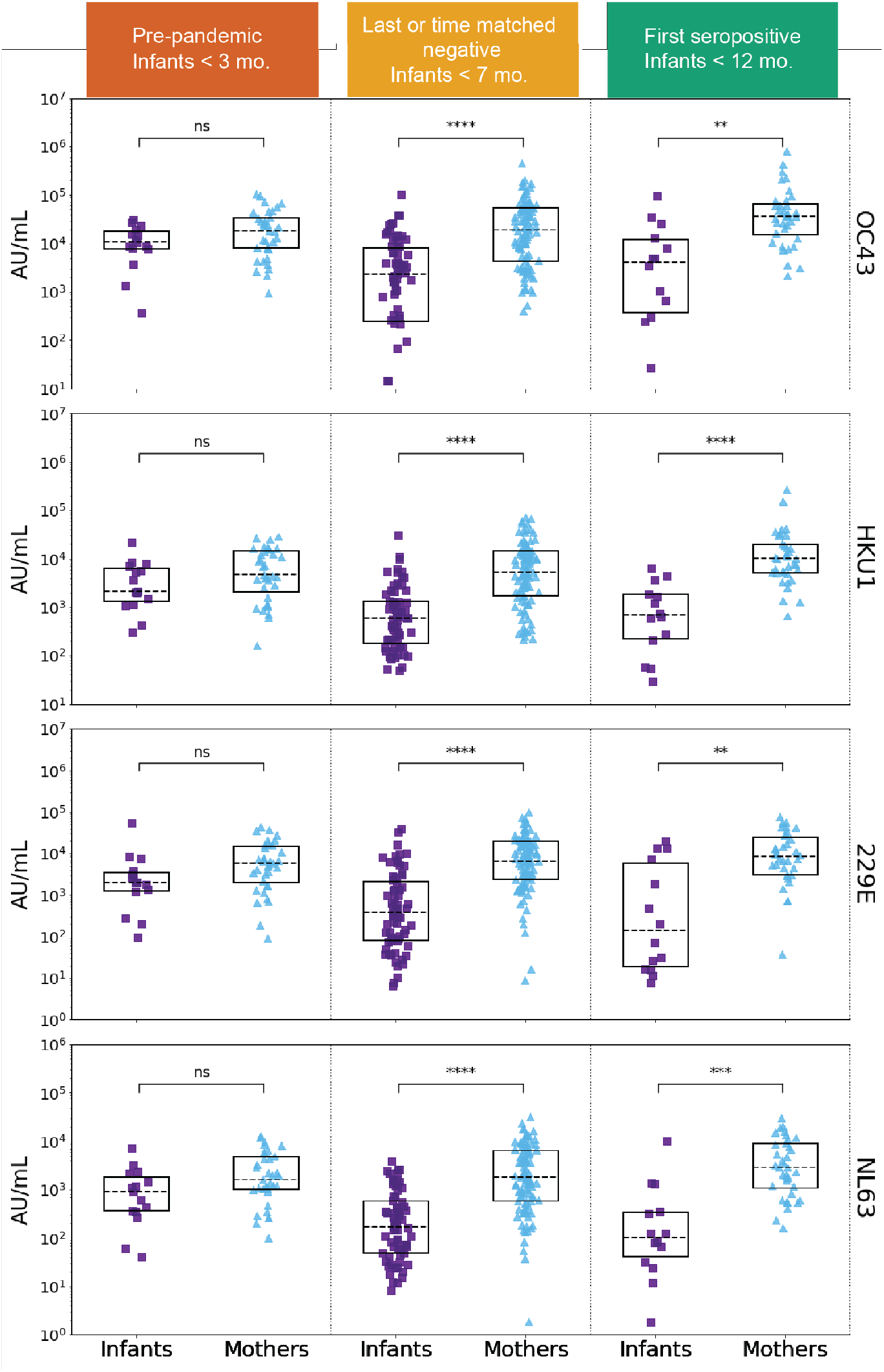
eHCoV IgG titers in SARS-CoV-2 naïve and SARS-CoV-2 seroconverted plasma from infants and mothers. HCoV-OC43, HCoV-HKU1, HCoV-229E, and HCoV-NL63 spike IgG levels (AU/mL) in pre-pandemic (SARS-CoV-2 naïve) plasma from never SARS-CoV-2 seropositive and eventually seroconverting infants (purple, N = 15) and mothers (blue, N = 35), last negative before SARS-CoV-2 seropositive or time-matched never seropositive infants (N = 67) and mothers (N = 97), and first SARS-CoV-2 seropositive samples from infants (N = 14) and mothers (N = 36). Sample groups and median infant age are indicated in the colored headings. P-values (Wilcoxon rank-sum test) are corrected for multiple hypothesis testing (Holm-Bonferroni). (ns) P > 0.05, (*) P ≤ 0.05, (**) P ≤ 0.01, (***) P ≤ 0.001, (****) P ≤ 0.0001.

### SARS-CoV-2 infection is associated with increases in betacoronavirus eHCoV antibody response

Our observation of significantly higher levels of eHCoV antibodies in mothers versus infants after SARS-CoV-2 seroconversion prompted us to test whether antibody levels increased between the last seronegative and first seropositive samples in infants and mothers. To test whether SARS-CoV-2 infection was associated with increases in antibodies that bind to eHCoV in infants and mothers, we compared antibody levels longitudinally in SARS-CoV-2 seroconverters. Between the last negative and first seropositive plasma samples, antibody levels for both endemic betacoronaviruses, HCoV-OC43 and HCoV-HKU1, increased significantly in mothers, but not infants, suggesting a cross-reactive response that could be influenced by pre-existing eHCoV antibodies in adults (**Figs. 3A, 3B and S1**). Alphacoronavirus antibody levels did not increase in either group. Interestingly, antibody levels against the highly pathogenic betacoronavirus, SARS-CoV-1, increased most significantly between the last negative and first seropositive samples, and this was true for both mothers and infants (**Figs. 3A, 3B and S1**). Given the lack of SARS-CoV-1 circulation, this result suggests a prior exposure to the virus is not driving this increase, rather it reflects *de novo* responses to SARS-CoV-2 infection that recognize SARS-CoV-1, which shares a high degree of sequence homology.

**Figure 3.**
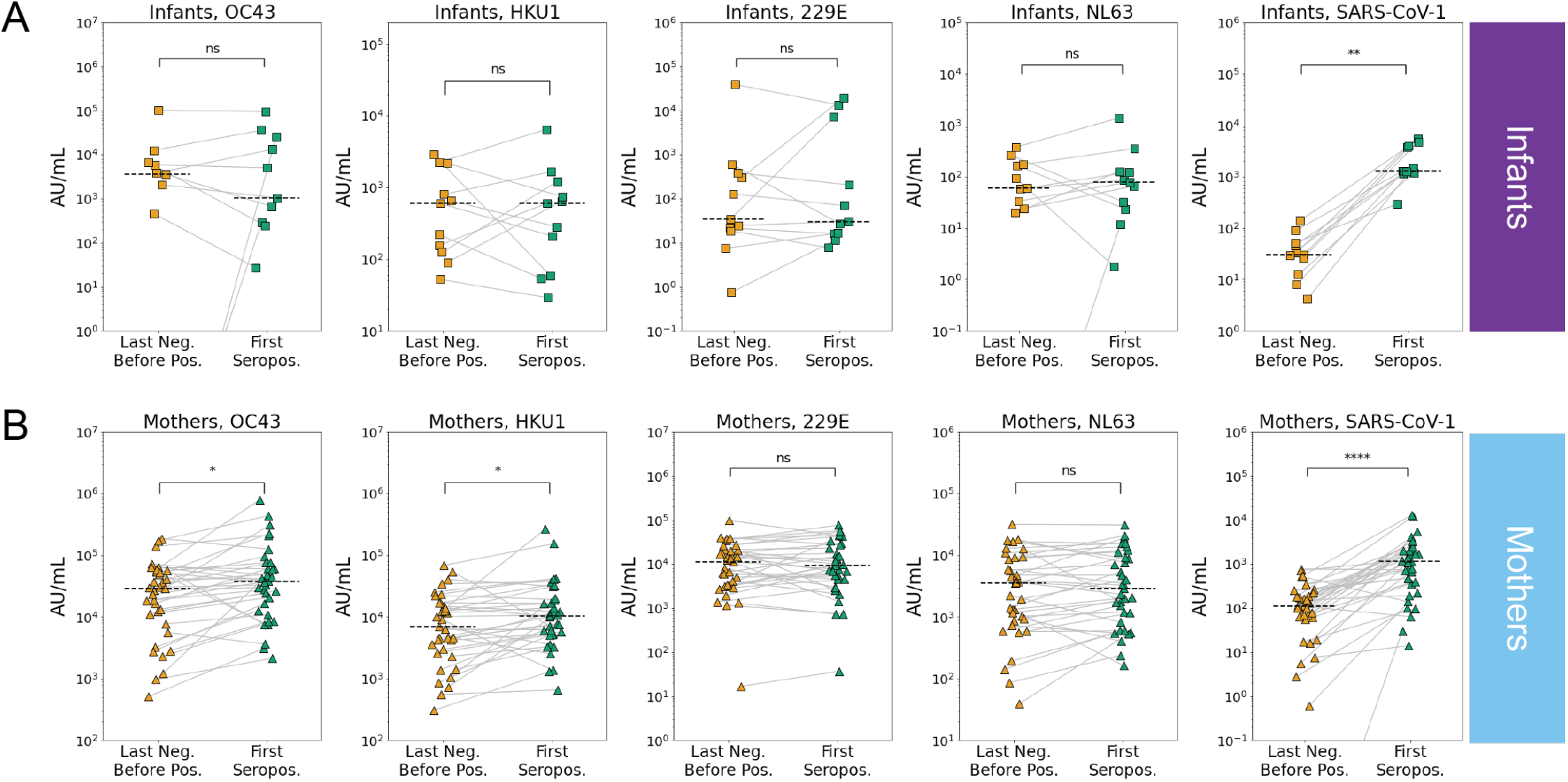
eHCoV and SARS-CoV-1 antibody titers immediately prior to and after SARS-CoV-2 seroconversion in infants and mothers. Last negative (yellow) and first seropositive (green) eHCoV spike IgG titers (AU/mL) in (A) infants (N = 11) and (B) mothers (N = 35). P-values (Wilcoxon signed rank test) are indicated and corrected for multiple hypothesis testing (Holm-Bonferroni). Significant comparisons (p < 0.05) are further indicated with an asterisk. (ns) P > 0.05, (*) P ≤ 0.05, (**) P ≤ 0.01, (***) P ≤ 0.001, (****) P ≤ 0.0001.

### Pre-existing eHCoV antibody levels are not strongly associated with SARS-CoV-2 serostatus

To test whether recent eHCoV antibody levels were associated with SARS-CoV-2 seroconversion, we compared eHCoV antibody binding titers between never seropositive and seroconverting infants and mothers in the last negative and time-matched seronegative samples. While we did not observe a relationship between prior eHCoV antibody titer and SARS-CoV-2 infection in mothers, we observed a modest association between HCoV-229E antibody binding titer and SARS-CoV-2 seronegativity in infants, though this result fell below statistical significance after correction for multiple hypothesis testing (**Figs. 4A and 4B**). Similarly, we did not observe a statistically significant relationship between pre-pandemic eHCoV antibody binding titer and SARS-CoV-2 seroconversion in infants or mothers (**Fig. S2**), though the number of samples in this group was more limited.

**Figure 4.**
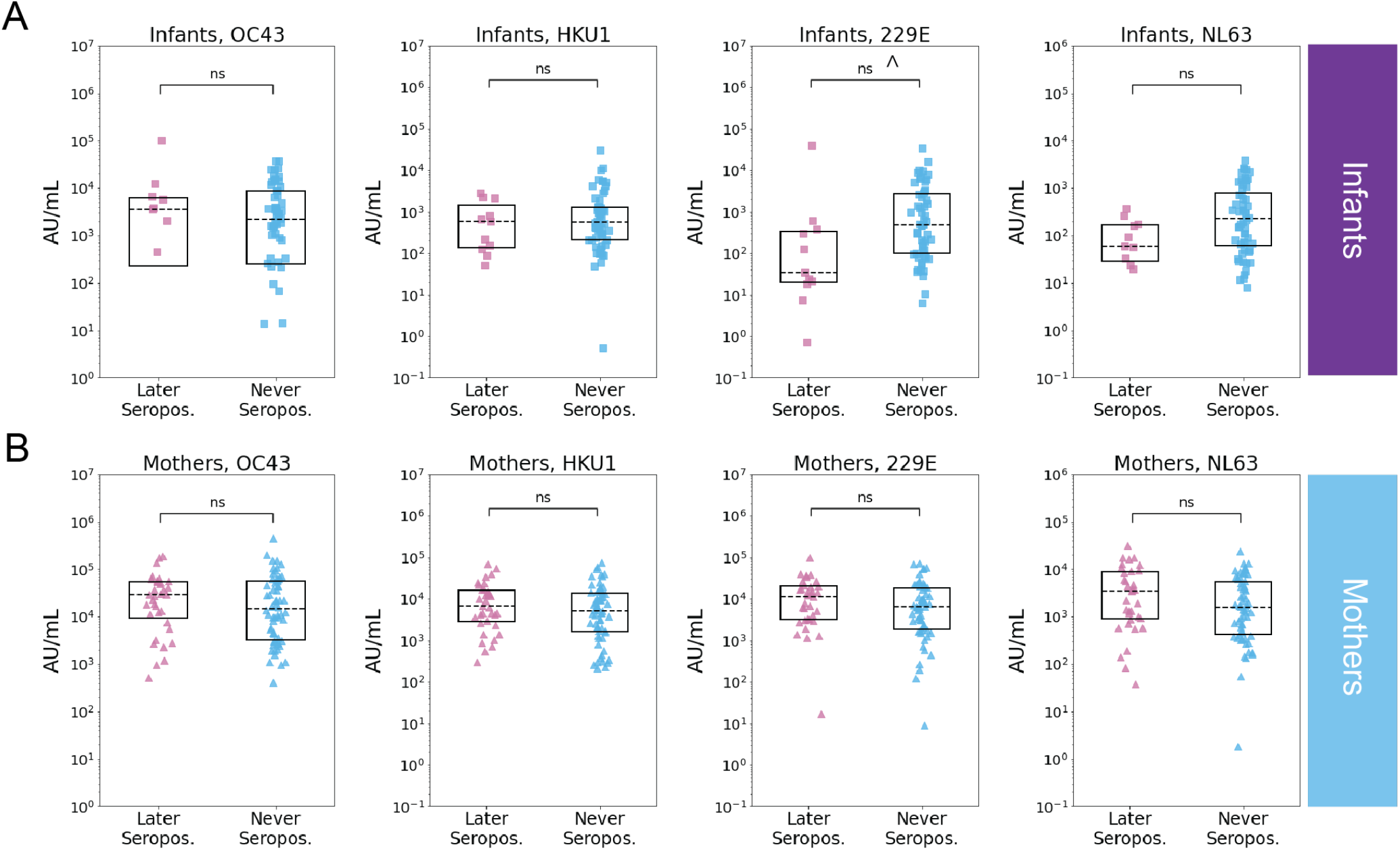
Relationship between last negative samples and SARS-CoV-2 serostatus in infants and mothers. Recent prior eHCoV spike IgG levels (AU/mL) in individuals that were later SARS-CoV-2 seropositive (pink) or seronegative (blue) in (A) infants (later seropositive N = 11, never seropositive N = 56) and (B) mothers (later seropositive N = 35, never seropositive N = 62). P-values (Wilcoxon rank-sum test) are indicated and corrected for multiple hypothesis testing (Holm-Bonferroni). (ns) P > 0.05, P ≤ 0.05 prior to Holm-Bonferroni correction indicated with ^.

## Discussion

The role of cross-reactive eHCoV antibody responses in SARS-CoV-2 infection or protection remains unclear and there is a scarcity of data on this relationship in the earliest months of life. In this study, we measured eHCoV IgG antibody binding responses in infants and mothers just prior to and after SARS-CoV-2 seroconversion. We found higher levels of eHCoV antibody binding in mothers versus infants just prior to SARS-CoV-2 infection, which likely reflects the decreased probability of exposure to eHCoVs during the shorter infant lifespan.

Increased eHCoV antibody levels upon SARS-CoV-2 seroconversion have been observed in some cohorts [3–10], but not others [18, 19], and, when detected, have been hypothesized to reflect boosting of pre-existing cross-reactive responses. We observed significant increases in betacoronavirus, but not alphacoronavirus, antibody levels (SARS-CoV-1 in infants and mothers; HCoV-OC43 and HCoV-HKU1 in mothers) upon SARS-CoV-2 seroconversion compared to the most recent seronegative sample, suggesting that there are cross-reactive antibodies to SARS-CoV-2, and they may be more likely to be present in the context of more closely related eHCoVs. Interestingly, we did not observe increased eHCoV antibody levels upon SARS-CoV-2 seroconversion in infants, which likely reflects the absence of pre-existing eHCoV memory responses in infants due to both passive antibody protection and a limited period of exposure. We identified SARS-CoV-1 cross-reactive antibodies in both infants and mothers, responses that are unlikely to reflect pre-existing memory responses given the lack of circulating SARS-CoV-1. Such cross-reactive responses suggest SARS-CoV-2 infection may induce naïvely generated cross-reactive responses that recognize SARS-CoV-1, though the above analyses did not model for time due to confounding factors including time since infection, postpartum date in mothers, and changing maternal antibody levels in infants.

Pre-existing immune cross-protection and lower median age have been hypothesized as correlates of protection against severe COVID-19 in Sub-Saharan Africa [20, 21]. However, whether pre-existing eHCoV antibodies are protective or increase risk of SARS-CoV-2 infection remains controversial [22], and this relationship is particularly understudied in infants. In our study, prior eHCoV antibody titer was not strongly associated with SARS-CoV-2 seroconversion. Further study with larger cohorts will be needed to evaluate this association, as our study is limited by sample size and potential heterogeneity in SARS-CoV-2 exposure risk in the study population. Together, these results demonstrate differences in eHCoV antibody responses pre-and post-SARS-CoV-2 infection between infants and mothers in Kenya, including evidence for HCoV-OC43 and HCoV-HKU1 antibody boosting upon SARS-CoV-2 seroconversion in mothers but not infants, and provide a basis for further evaluation of cross-reactive eHCoV antibody responses in newborns and young infants in the context of SARS-CoV-2.

## Materials and Methods

### Study participants

A subset of mothers and infants in Nairobi, Kenya that were already enrolled in the Linda Kizazi Study, a prospective cohort of mother-to-child virome transmission, consented to SARS-CoV-2 serology testing as previously described (Begnel et al., in revision). Mother-infant pairs attended clinic visits approximately every 3 months, at which time clinical data was collected including recent diagnoses and healthcare visits, symptoms of illness at the visit or since the last visit, and history of current or recent medications or immunizations. Physical exams were conducted at each clinic visit and samples including blood were collected. The Kenyatta National Hospital-University of Nairobi Ethics and Research Committee and the University of Washington and Fred Hutchinson Institutional Review Boards approved of all Human Subjects study procedures.

### Sample Selection

Plasma samples collected between April 2019-December 2020 were selected for this sub-study based on previous SARS-CoV-2 serology testing (Begnel et al., in revision). For participants that seroconverted during the study period, up to 3 longitudinal plasma samples were included: the “first seropositive” was the plasma sample in which SARS-CoV-2 antibodies were first detected by ELISA testing (Begnel et al., in revision), the “last negative” was the most recent plasma sample collected prior to the first seropositive sample time point, and a “pre-pandemic” sample (if available) was the most recent sample collected prior to October 2019 to ensure no possible exposure to SARS-CoV-2. Samples from participants that did not seroconvert to SARS-CoV-2 during the study period include up to 2 longitudinal plasma samples: a “time-matched seronegative” sample, which was collected during the time window of the last negative sample from the seroconverters (December 2019-April 2020, and a “pre-pandemic” sample, as described above.

### Multiplexed chemiluminescent antibody binding assay with plasma

Plasma samples were heat-inactivated for 60 minutes at 56°C prior to 1:5000 dilution. eHCoV antibody levels were determined using Mesoscale Diagnostic’s V-PLEX Coronavirus Panel 2 which includes all four eHCoVs, plus SARS-CoV-1 spotted together in individual wells in a 96-well format. The assay was performed following the manufacturer’s instructions. Diluted samples, along with manufacturer-provided calibrators and controls, were applied to blocked plates and incubated for 2 hours. Washed plates were incubated with detection antibody for 1 hour, followed by addition of MSD GOLD Read Buffer B. Plates were read on MSD instrument and raw data were processed in MSD Discovery Workbench software (version 4.0). IgG antibody levels for each antigen were calculated in Workbench based on the calibrator standard curve fit and reported in Arbitrary Units/mL (AU/mL).

### Statistical analyses

Wilcoxon rank-sum or Wilcoxon signed-rank tests were performed to compare samples that were unmatched or matched, respectively. P-values were adjusted for multiple hypothesis testing by applying Holm-Bonferroni correction. All statistical tests were performed using SciPy [23] and statsmodels [24] software tools.

## Acknowledgements

We thank all study volunteers for their participation. We thank Carolyn Fish for assistance with sample preparation. We are grateful to the Linda Kizazi study team and members of the Overbaugh and Lehman labs for helpful discussion and advice. This work was funded by Operating Grants from the Canadian Institutes of Health Research (PI Gantt), and National Institutes of Health grants HD092311 (PI Lehman) and AI138709 (PI Overbaugh).

---

**Figure S1.**
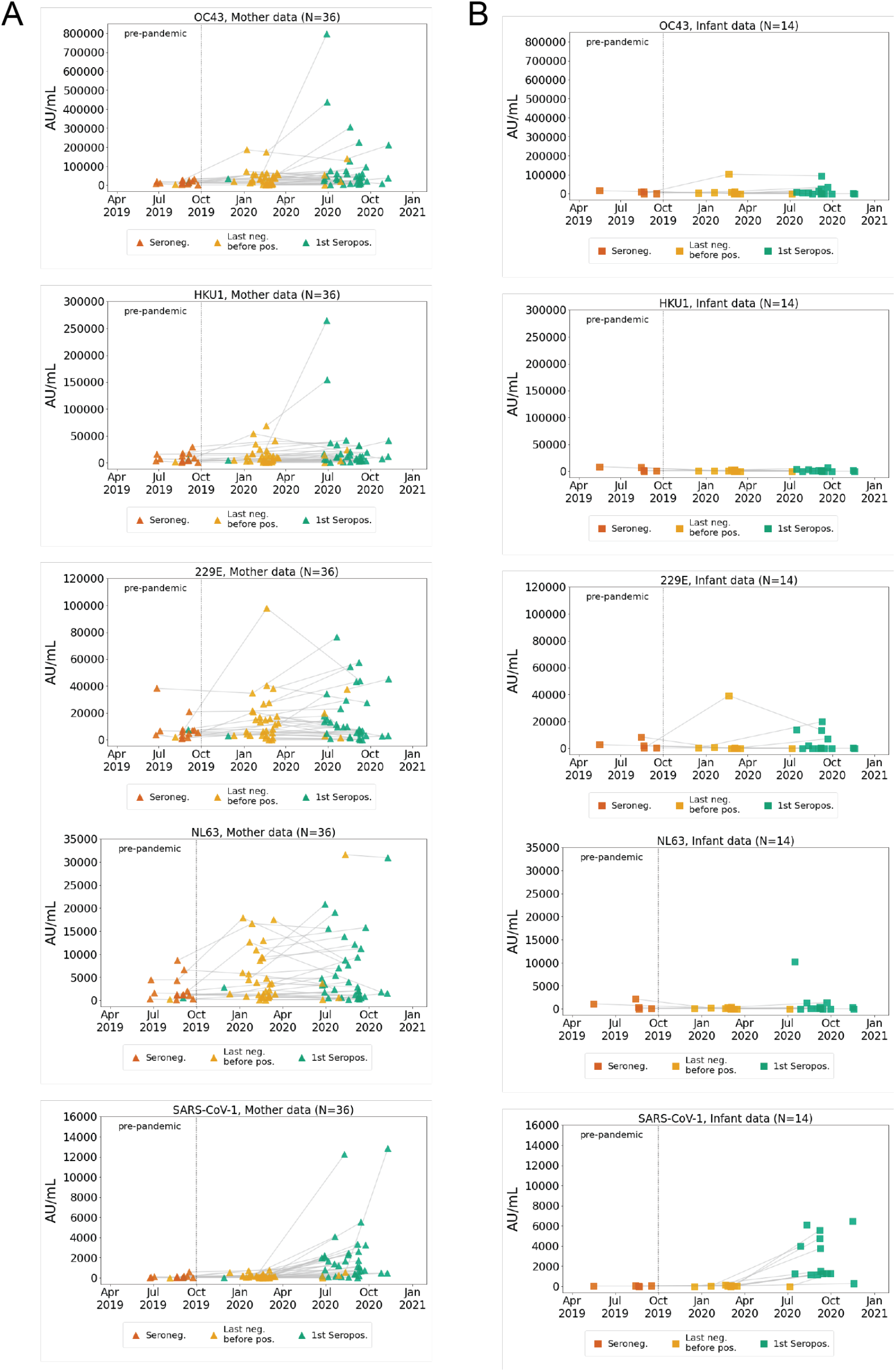
Longitudinal antibody binding responses to eHCoVs and SARS-CoV-1 in individuals that eventually seroconverted to SARS-CoV-2. IgG titers (AU/mL) over time for indicated HCoVs; left panels, eventually SARS-CoV-2 seroconverting mothers (N = 36); right panels, eventually SARS-CoV-2 seroconverting infants (N = 14).

**Figure S2.**
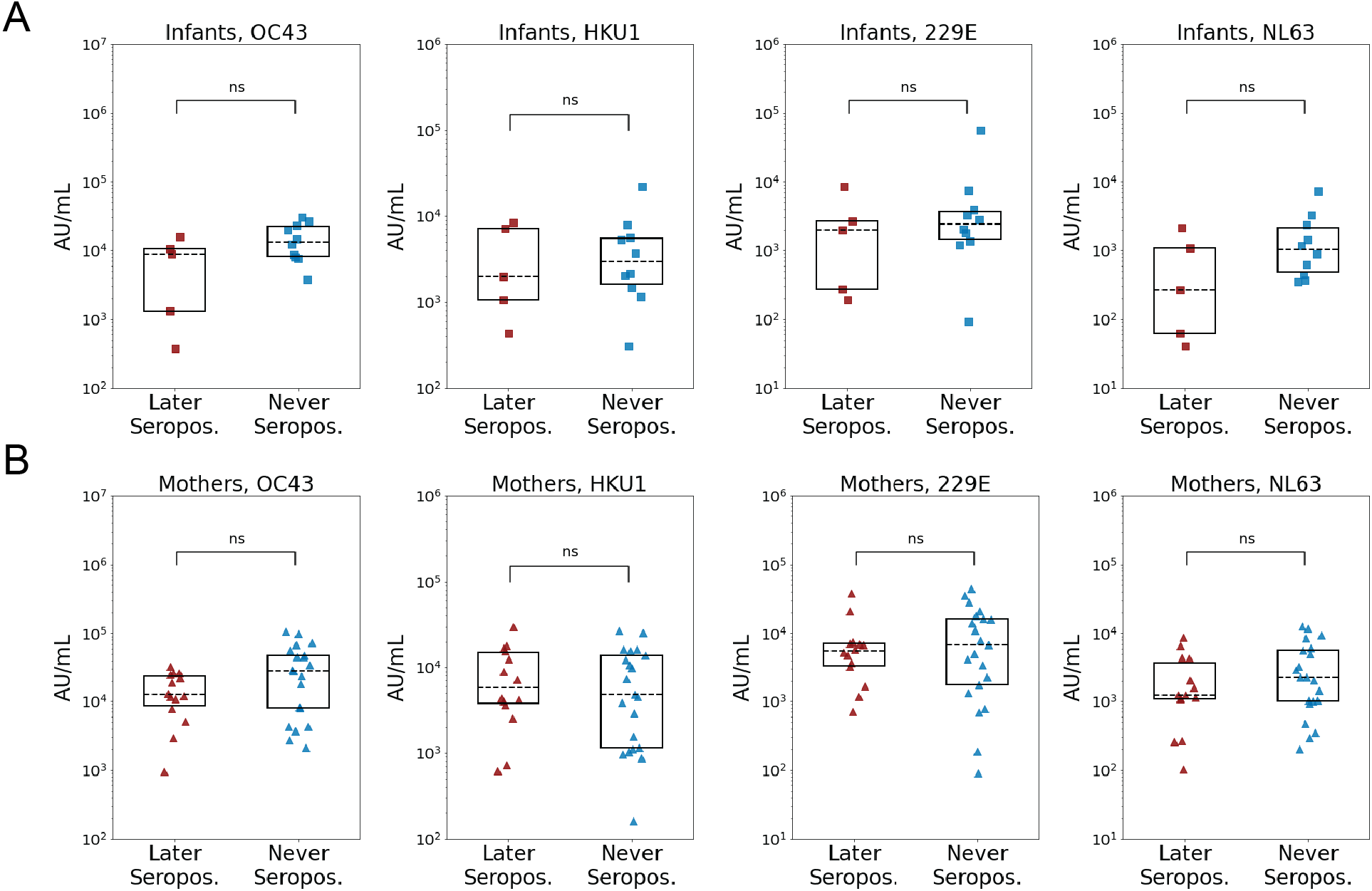
Relationship between pre-pandemic eHCoV antibody titer and SARS-CoV-2 serostatus in infants and mothers. (A) eHCoV antibody titers in infants that later became seropositive (N = 5) or were never seropositive (N = 10) for SARS-CoV-2. (B) eHCoV antibody titers in mothers that later became seropositive (N = 14) or were never seropositive (N = 21) for SARS-CoV-2. P values (A and B) were calculated using Wilcoxon rank-sum test with Bonferroni correction for multiple hypothesis testing. (ns) P > 0.05, (*) P ≤ 0.05, (**) P ≤ 0.01, (***) P ≤ 0.001, (****) P ≤ 0.0001.

